# Developmental emergence of quiescent-like neural progenitor cells in the zebrafish embryonic brain

**DOI:** 10.1101/2024.02.13.580083

**Authors:** Yuanyuan Liu, Carla-Sophie Lembke, Judith TML Paridaen

## Abstract

The mature brain is made up of differentiated neurons and glial cells that are produced by embryonic neural stem and progenitor cells (collectively called neural stem cells (NSCs)) during brain development. In contrast, adult NSCs generate only a limited number of neuronal and glial cell types. Moreover, in contrast to embryonic NSCs, adult NSCs spend most of their time in a non-dividing resting phase called quiescence. In the adult brain, the quiescent NSCs are activated at a low frequency to divide and produce new cells to replace lost or damaged mature cells.

Previous studies have demonstrated that about halfway through mammalian brain development, a subset of the embryonic NSCs population slows down their division rate and slowly transition into adult quiescent NSCs (qNSCs). However, the molecular mechanisms that underlie the emergence of the pre-adult SCs remain largely unknown.

Here, we explored single-cell transcriptomes from several embryonic stages of zebrafish development in order to determine at what developmental stage transcriptional signatures typical of adult quiescent NSCs first emerge. We identified a subpopulation of embryonic NSCs with a distinct transcriptional profile from other embryonic NSCs. This population shares transcriptional similarities with adult qNSCs including genes known to maintain quiescence. We propose that this population constitute slower cycling embryonic NSCs that may transition into the adult qNSCs of the adult zebrafish brain.

## Introduction

Neural stem and progenitor cells (NSCs) play a key role in the intricate processes of embryonic development and adult brain homeostasis. These cells are the primary architects behind the generation of neurons during the early stages of embryonic growth. Moreover, in vertebrates, limited numbers of adult neural stem cells support life-long generation of new neurons. While embryonic NSCs divide continuously to facilitate brain growth, adult NSCs spend most of their time in a non-dividing resting phase called quiescence (Urbán and Cheung, 2021; Velthoven and Rando, 2019). Quiescence is defined as a reversible cell cycle state where cells are temporarily arrested in G0 phase. Here, quiescent adult NSCs are activated at a low frequency to divide (Obernier and Alvarez-Buylla, 2019). Once activated, adult neural SCs generate limited numbers of new cells such as glial cells and neurons (Obernier and Alvarez-Buylla, 2019; Youssef et al., 2018).

In the developing mammalian brain, a subset of the embryonic NSC population slows down their division rate and obtains a pre-adult neural SCs identity (Berg et al., 2019; Falk et al., 2017; Fuentealba et al., 2015; Furutachi et al., 2015). In the developing mouse forebrain, the CKI p57 is required for induction of slower cell cycles in early pre-adult neural SCs(Furutachi et al., 2015), suggesting that expression of CKIs induces quiescence *in vivo*. Later in life, these slow-cycling cells transition into the adult quiescent NSCs (qNSCs). Recent work in mice showed that late embryonic NSCs share gene transcriptional signatures with adult quiescent NSCs, showing that there is a gradual transition (Yuzwa et al., 2017). However, there is little known about the molecular mechanisms that underlie the specification of a subset of embryonic NSCs as pre-adult NSCs (Hu et al., 2017; Than-Trong et al., 2018). It is also unclear whether pre-adult NSCs stem from random sampling or specific selection based on certain characteristics within embryonic NSCs (Berg et al., 2019).

The zebrafish constitute a popular model organism in neurobiology, as they, in contrast to mammalian systems, show widespread adult neurogenesis over many regions of the brain (Kizil et al., 2012; Labusch et al., 2020). Moreover, they possess the remarkable ability to regenerate neurons following an injury. The characteristics of zebrafish embryonic and adult NSCs are largely similar to those of their mammalian counterparts (Jurisch-Yaksi et al., n.d.; Labusch et al., 2020). However, there is very little information regarding the developmental emergence of pre-adult NSCs. Previous work has indicated that in the zebrafish developing telencephalon, the proportion of proliferating embryonic NSC is decreasing between 5 and 10 days-post-fertilization (dpf; (Alunni et al., 2013; Than-Trong et al., 2018). This suggests that, similarly to mammalian systems, a subset of embryonic NSC slows down their cell cycle and enters quiescence as the pre-adult NSCs at late stages of brain development.

Here, we made use of a recently published single-cell transcriptomics dataset covering the developing zebrafish brain from 12 hours to 15 days post-fertilization (Raj et al., 2020). We specifically investigated the expression profile and changes therein of embryonic NSCs, and examined whether we could find sub-clusters of NSCs with gene expression profiles related to quiescence. We found that a subset of embryonic NSCs showed a gene expression signature similar to adult quiescent NSCs from 3 dpf onwards. This suggests that the specification of pre-adult NSCs in the zebrafish brain occurs earlier than previously proposed.

## Results

### Single-cell profiling and unbiased clustering of zebrafish neural progenitor cells

The primary objective of this investigation was to assess whether transcriptional signatures of pre-quiescent NSCs can be identified at zebrafish embryonic stages. As input, we used a previously published comprehensive single-cell atlas of zebrafish brain development comprising 12 developmental stages between 12 hpf and 15 dpf (Raj et al., 2020) (Figure 1A). To assess the distinctions among neural progenitor cells and to avoid potential confounding effects from other cell types, particularly neurons, we included only cells previously annotated as progenitor cells and cycling cells in our analysis.

**Figure 1.**
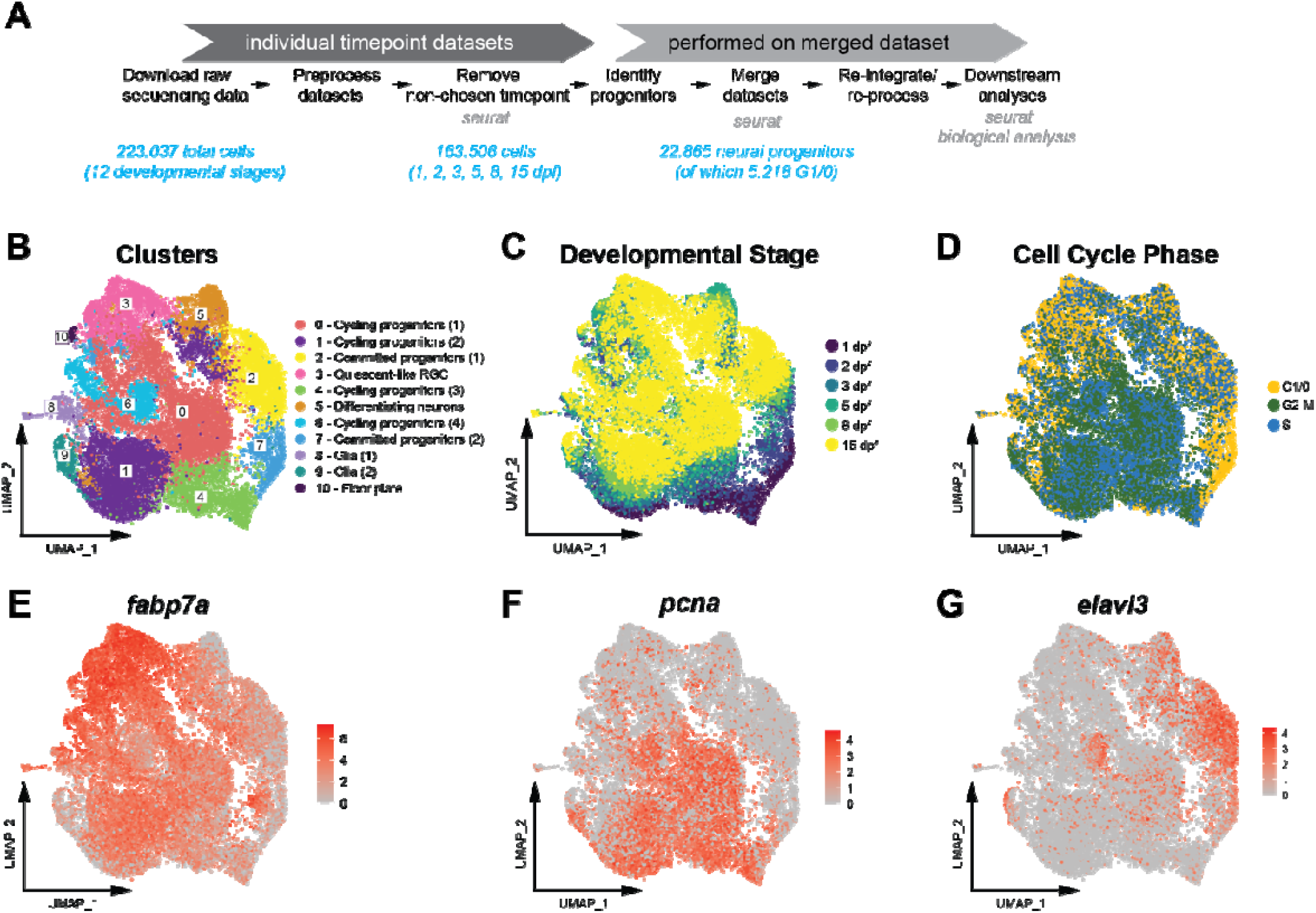
Progenitor cell composition of the embryonic zebrafish brain between 1 and 15 dpf. (A) Overview of the steps within the analysis pipeline with the number of cells included in blue. (B) Unsupervised UMAP plot containing all 22.865 neural progenitors from 6 stages that separate into 11 distinct clusters. (C) Projection of the developmental stage of each individual cell onto the UMAP plot. (D) Projection of the predicted cell cycle phase of each individual cell onto the UMAP plot. (E-G) UMAP plots showing the log-normalized expression levels for the cell type marker genes fabp7a (radial glial cells; (E), pcna (proliferating cells; (F) and elavl3 (differentiating neurons; (G)). The expression level color scales and cluster identification keys are provided next to the plots.

The bioinformatics workflow, delineated in Figure 1A, was systematically stratified into two discernible phases: processes executed on individual developmental stage datapoints and those applied to the merged dataset. We included all stages, except the 36 hpf timepoint, between 1 and 15 dpf in our analysis. Within the dataset containing the selected developmental stages comprising of 163,506 cells, we filtered out all cells designated as progenitor cells, a process that was crucial in setting the stage for our primary objective. Since we expected that a potential qNSC-like cluster would be composed of very few cells in comparison to the total cell population within the developing brain, including a large variety of other cell types would only confound the smaller differences present within the progenitor population. Filtering out all non-progenitor cells yielded 22,865 progenitor cells across all the timepoints.

Next, we applied Seurat to the merged dataset of 22.865 progenitor cells to cluster together transcriptionally similar cells. After testing and selecting an appropriate clustering resolution value (Figure S1A), we obtained 11 distinct clusters as shown by the Uniform Manifold Approximation and Projection (UMAP) plot (Figure 1B). When exploring the clusters in more detail according to the developmental stage of the individual cells (Figure 1C) we observed that none of the clusters separated solely based on their timepoint of developmental origin. Instead, each cluster was comprised of cells originating from multiple timepoints. This suggests a transcriptional continuity in cell type identity that connects cells across multiple developmental stages.

To assign the identity of each cluster, we compared the most enriched genes (Figure S2) with the cell type annotations of the original dataset and other typical cell type marker genes (Figure S3). Based on the cell type markers (Figure S2) and expression of typical neural progenitor markers such as *fabp7a* (Figure 1E, Figure S4E), it appeared that most of the cells indeed constitute neural progenitor cells. Together, this initial analysis shows that amongst the population of NPCs at embryonic and larval stages, separate subclusters with distinct transcriptomes emerge that may be indicative of biological differences.

### Classification of NPCs based on cell-type specific marker genes

To obtain more insights into the nature of the different cell clusters, we next explored the cell cycle stage of each cluster. Clusters 0, 1, 4, 6 and 9 showed clear enrichment of cells in G2-M and S-phase of the cell cycle (Figure 1D, Figure S1B). Moreover, many of the cells in these clusters express the proliferating cell marker *pcna* (Figure 1F). Therefore, we assigned cycling neural progenitor identity to these clusters.

Several clusters (3, 7, 10) demonstrated an enrichment of individual cells in G1/0 phase (Figure 1D, Figure S1B). Moreover, clusters 2, 3, 5, 7, 8 and 10 showed a relatively smaller proportion (<25% of total) of cells in G2-M phase. This suggests that these NPCs may progress through the cell cycle more slowly than the other NPC clusters. This could result from these cells being committed to differentiation (which is linked to slower cell cycle progression (Calegari et al., 2005; Lange et al., 2009). Alternatively, they may constitute slower cycling uncommitted NPCs, such as pre-quiescent NSCs (Furutachi et al., 2015).

Exploration of the top enriched genes in the clusters with lower proportions of S and G2-M phase showed that cluster 7 expressed known proneural genes such as *atoh1c* and *neurod1* (Figure S2). Clusters 2 and 5 also expressed neuronal markers such as *elavl3* and *elavl4* (Figure 1G, Figure S1C). This suggests that clusters 2, 5 and 7 constitute NPCs that are neurogenic (committed to neuronal differentiation).

To explore whether any of the clusters could constitute pre-quiescent NSCs, we assessed the expression of known zebrafish adult qNSCs markers (Lange et al., 2020; Than-Trong et al., 2018) in the merged dataset. We found that cluster 3 showed consistent enrichment of many adult qNSC cell markers such as *glula, cx43* and *mfge8a* (Figure S2 and S4E; (Lange et al., 2020)). This suggests that cluster 3 constitute pre-quiescent slower cycling NPCs.

Since we expect pre-quiescent NPCs to be mainly in G1/0 cell cycle stage, we also performed unsupervised Seurat clustering for G1/0 only cells (Figure S4C, D, F). We found 8 separate clusters with a similar distribution over the developmental timepoints as the allNPCs dataset (Figure S4A-D). Exploration of qNSC marker gene expression showed one cluster (0) with consistent enrichment of qNSC marker genes (Figure S4F). This confirms that there appears to be a subpopulation of embryonic G1/0 NPCs (qNSC-like NPCs) with transcriptional signatures similar to adult qNSCs.

### Emergence of qNSC-like NPC cluster during embryonic stages

To gain further understanding in the emergence of the qNSC-like NPCs during development, we next returned to the individual timepoints, ran the Seurat clustering (Figure S5) and explored the transcriptional signatures of each cluster, looking specifically at the adult qNSC marker genes at each developmental timepoint (Figure 2).

**Figure 2.**
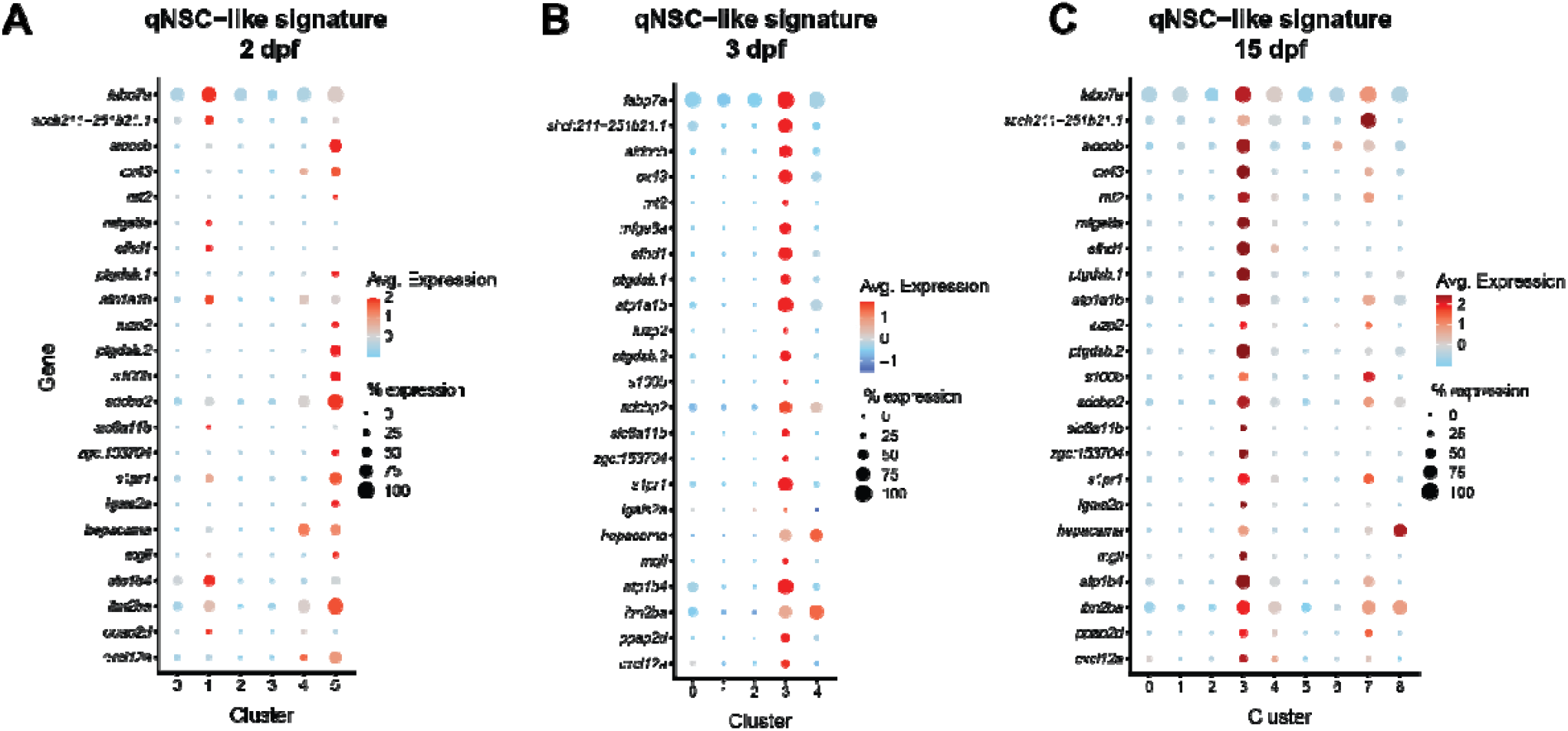
The presence of qNSC-like transcriptional signatures in neural progenitor clusters of the embryonic and larval zebrafish brain. Dotplots showing the gene expression of adult qNSC markers at (A) 2 dpf, (B) 3 dpf, and (C) 15 dpf. The y axis shows the individual marker genes and the x axis shows the progenitor clusters at each timepoint. The size of the dots reflects which percentage of cells within a given cluster which express the gene in question while the color of the dots represents the log-normalized expression level with red signifying high expression and blue low expression as shown in the individual keys.

At 1 dpf, none of the clusters exhibited genes expressing positive signatures for these markers (Figure S1D). By 2 dpf, cluster 5 displayed enrichment for some of the adult qNSC marker genes (Figure 2A). By 3 dpf, cluster 3 exhibited a robust and distinct signature for the majority of the adult qNSC marker genes (Figure 2B). Analysis of the subsequent time points consistently revealed one NPC cluster with expression profiles aligning with the adult qNSC gene markers. Specifically, at 5 dpf, cluster 4 exhibited the most pronounced expression of adult qNSC markers, while cluster 7 demonstrated a weaker but analogous gene expression profile (Figure S1E). At 8 dpf, cluster 3 exhibited the highest expression of adult qNSC markers, and cluster 6 displayed a weak expression profile that partially overlapped with adult qNSCs (Figure S1F). Finally, at 15 dpf, cluster 3 emerged as the most representative of cells with high expression levels of the marker genes, with cluster 7 demonstrating moderate gene expression levels of some of the adult qNSC marker genes (Figure 2C).

Our analysis encompassed 23 marker genes stemming from earlier studies on adult qNSCs in the zebrafish brain (Lange et al., 2020; Than-Trong et al., 2018). Most genes exhibited expression in a subset of NPCs for at least one timepoint. Notably, *luzp2, s100b*, and *lgals2a* consistently demonstrated absence of expression enrichment in the qNSC-like and other clusters. The remaining 20 adult qRGC marker genes exhibited expression in qNSCs-like clusters at one or more time points. Furthermore, 16 marker genes consistently delineated clear qRGC-like clusters across all temporal stages from 3 dpf.

Together, the dynamics of the expression levels of qNSC-marker genes shows that a subset of embryonic NPCs with an expression profile resembling that of adult qNCS emerges at 3 dpf of zebrafish development.

### Quiescent-like embryonic neural progenitors express core identity genes throughout development

Genes that are transcriptionally restricted to qNSC-like clusters and that are consistently expressed in these clusters may represent genes that promote the establishment and maintenance of qRGCs and exemplify their core transcriptional identity. Therefore, having identified a subpopulation of quiescent-like progenitors within the general embryonic NPC population, we next performed a comprehensive investigation into which genes were consistently enriched within this population. The overarching aim was to ascertain whether qNSC-like cells express specific genes early on or throughout the entirety of the developmental process that could play a role in their emergence.

First, we compared the transcriptomes of the qNSC-like cell clusters between individual timepoints to identify the significantly differentially expressed genes (DEGs) for each timepoint (Figure 3A). This resulted in a range of 244 (8 dpf) to 557 (15 dpf) DEGs. Next, we determined the overlap in the specific DEGs between timepoints. Through iterative comparisons of the DEGs in qNSC-like clusters relative to all other clusters between the developmental time points in which we found a robust qNSC-like cluster (3, 5, 8, and 15 dpf), a subset of DEGs specific to qNSC-like cell clusters was discerned (Figure 3B). Among the 1,710 DEGs identified in qNSC-like clusters across the four temporal stages, 123 exhibited consistent enrichment throughout the entire developmental continuum from 3 dpf onwards (Figure 3B).

**Figure 3.**
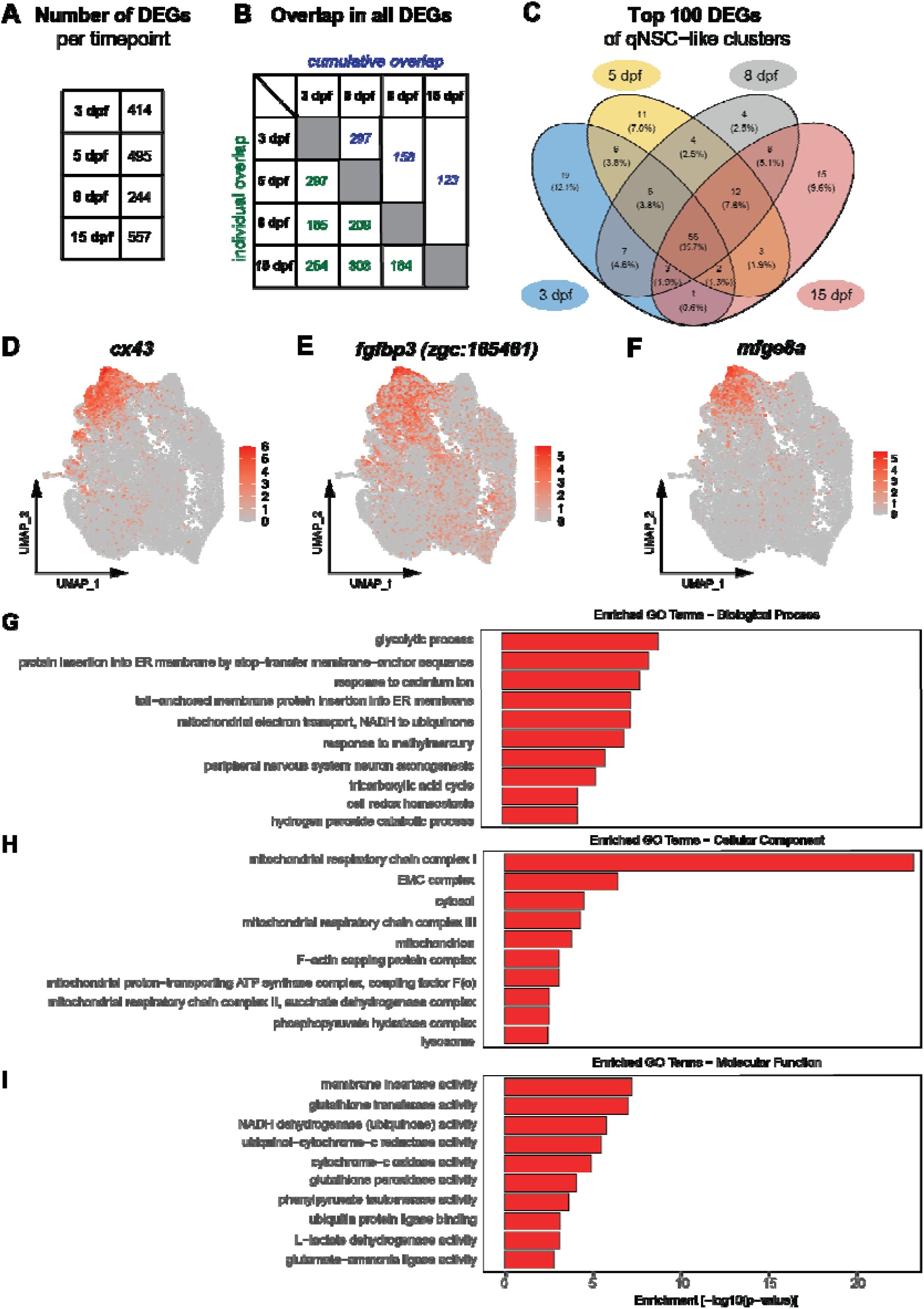
Quiescent-like neural progenitors express core identity genes throughout embryonic development. (A-C) Overview of overlapping DEGs across timepoints. (A) The number of DEGs in each of the qNSC-like clusters for each timepoint in which they were identified. (B) Quantification of the overlap in all DEGs of qNSC-like clusters across timepoints. The green values below the diagonal represent individual comparisons between timepoints, while the blue values above the diagonal represent cumulative comparisons across timepoints. (C) Venn-diagram showing the number of overlapping genes (and the relative proportion of the overlapping genes) between the individual stages (3 to 15 dpf). (D-F) UMAP plots showing the log-normalized expression levels for the qNSC marker genes *cx43* (D), *fgfbp3* (E) and *mfge8a* (F). The expression level color scales are provided next to the plots. (G-I) Enrichment scores of top 10 significantly enriched GO-terms for Biological process (G), Cellular component (H) and Molecular function (I) in the qNSC-like cluster. The bar position along the x-axis denotes the enrichment score that was calculated by taking the negative decadic logarithm of the p-value calculated with the Fisher’s test. GO-terms are listed on the y-axis.

Further refinement of the analysis focused on the DEGs with the highest expression levels (top 100 DEGs) at each developmental stage. Consequently, a catalogue comprising 56 genes emerged, characterized by heightened enrichment within each qNSC-like gene cluster across the entirety of embryonic development (Figure 3C). These genes can be broadly categorized into two groups. The first group encompasses genes previously acknowledged as markers for quiescent NSCs, including but not limited to *mfge8a, slc6a11b, cx43, atp1b4, s1pr1, atp1a1b, hepacama, ptgdsb*.*2, ptgdsb*.*1, aldocb*, and *efhd1*. The second group comprises genes whose characterization within the context of quiescence remains incomplete. This group includes *aqp1a*.*1, slc6a1b, cdocb, efhd1, slc6a1b, cxcl14, cyp26b1, slc38a3a, slc1a2b, zgc:165461 (fgfbp3), slc3a2a, ptn, slc4a4a, cd63, glula, abi3a, eno1b, mdkb, cyp2ad3, si:ch211-66e2*.*5, spry2, sulf2a, ndrg3a, cspg5b, smox*, and *id1*.*1*.

Exploration of the expression levels of *cx43, zgc:165461* (*fgfbp3*) and *mfge8a* specifically by mapping their expression levels on the UMAP plot revealed their restricted expression to mainly the qNSC-like cluster (Figure 3D-F). To derive more information regarding the potential function of the embryonic qNSC-like enriched genes, we conducted Gene Ontology (GO) term enrichment analysis of the qNSC-like cluster (Figure 3G-I). The analysis unveiled enrichment of terms pertaining to metabolic pathways (glycolysis, TCA cycle, mitochondria respiration) and transmembrane ion transport (Figure 3G-I).

Taken together, we identified a set of genes that are consistently expressed by embryonic qNSC-like progenitors, which may include genes involved in their initial emergence.

### Trajectory inference reveals diverging paths towards a neurogenic or qNSC-like identity

Next, we assessed the differentiation program at the transcriptional level as embryonic NPC transition into neurogenic and quiescent-like progenitor cells. Using Monocle3 trajectory inference, we systematically ordered cells from the 11 principal cell clusters based on their pseudotime, delineating their progressive differentiation states (Figure 4, Figure S6). We took the 1 dpf embryonic NPC cluster (4) as the starting point and root for the pseudotime ordering (Figure 4A). Two trajectories were revealed, one progressing towards a committed/neurogenic identity (right), and another one following cycling progenitors with a branching point towards the qNSC-like identity (left).

**Figure 4.**
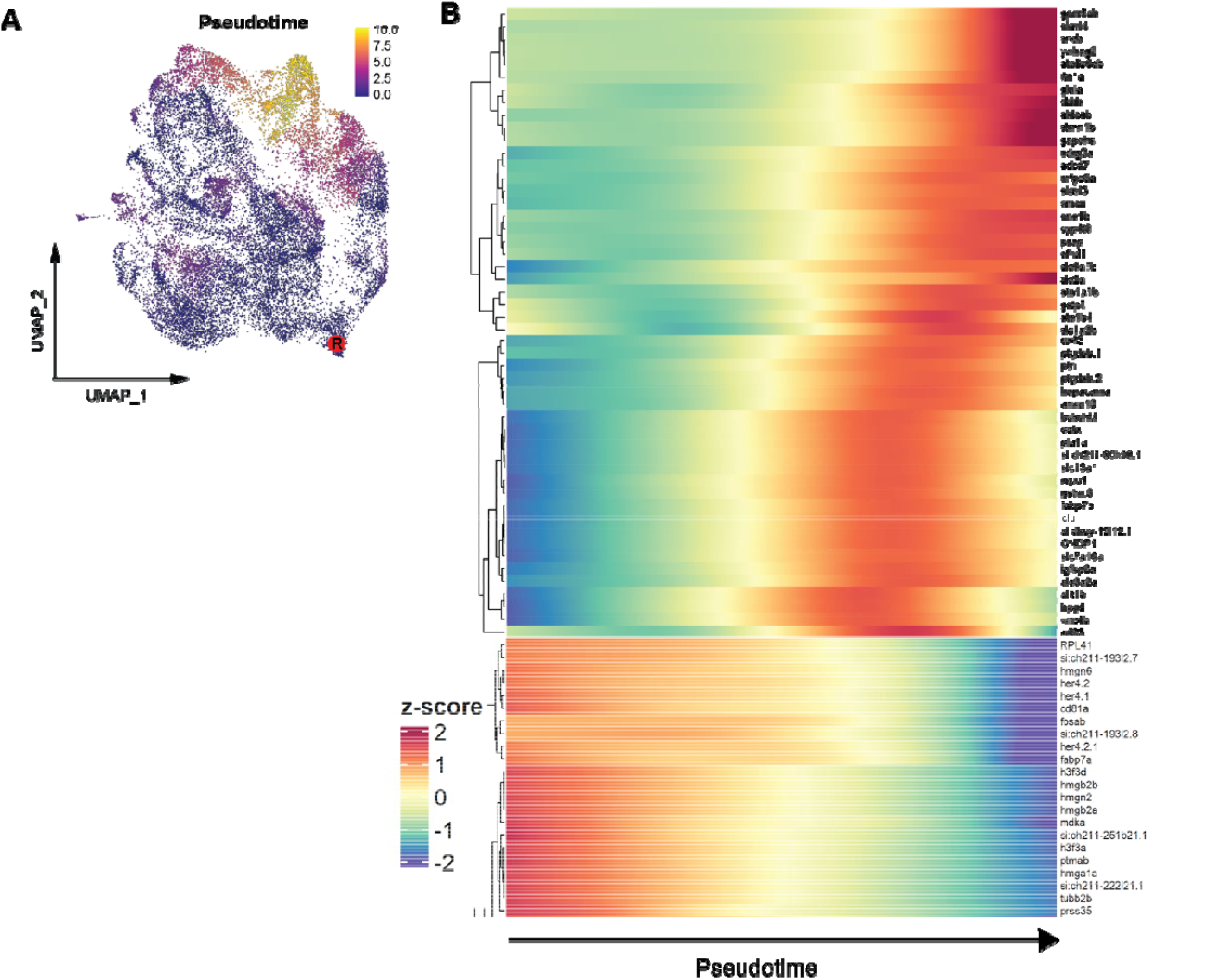
Pseudotime trajectory inference reveals diverging paths towards a neurogenic or qNSC-like identity. (A) Pseudotime assignments mapped to the UMAP visualization plot. R shows the root point used to calculate the pseudotime trajectory. Ordering in pseudotime (color key from early (0.0) to late 10.0)) reflects the predicted developmental trajectory. Two trajectories are revealed, one progressing towards a neurogenic identity (right), and another one following cycling progenitors with a branching point towards a qNSC-like identity (left). (B) Heatmap showing dynamic changes in expression levels for selected genes over pseudotime. Genes of interest include genes related to neurogenesis (*elavl3, elavl4*) and candidate genes related to quiescent-like state (*mfge8a, cx43*) and the radial glial marker gene *fabp7a*. The pseudotime z-scores are shown in the color keys.

Investigation of the genes with a dynamic expression pattern over pseudotime showed that there were several distinct expression patterns. In one pattern, gene expression levels decrease over pseudotime (Figure 4B, Figure S6). Many of the genes showing this pattern included typical NPC genes such as *her4.2* as well as genes encoding ribosomal proteins. Conversely, there were genes that were initially not expressed, but started getting expressed halfway or late through pseudotime. These genes included the adult qNSC marker genes *cx43, mfge8a* as well as the neuronal differentiation genes *elavl3* and *elavl4*. This suggests that there is a gradual transcriptional transition towards either differentiated or quiescent-like status.

Together, the pseudotime analysis confirms a gradual transcriptional trajectory of embryonic proliferating NPCs to committed/neurogenic NPCs and qNSC-like NPCs.

## Discussion

Here, we analyzed a previously published scRNAseq dataset to identify possible quiescent-like embryonic NPCs in the developing zebrafish brain. We found several subpopulations of embryonic NPCs and identified one particular subgroup of NPCs with a transcriptional signature similar to adult zebrafish qNSCs. We found that this signature emerged at 3 dpf and lasted up til at least 15 dpf, the latest timepoint in the dataset. The transcriptome of this quiescent-like embryonic NPC population contained a core set of genes that were expressed throughout this developmental period. This core set included known markers of adult qNSCs including *fabp7a, cx43, mfge8a* and *slc1a2b* (Dulken et al., 2017; Lange et al., 2020; Urbán et al., 2019).

Additional information that supports the conclusion that this NPC subpopulation constitutes quiescent-like or slow-cycling NPCs were the low expression levels of the proliferation marker *pcna*, the relatively high abundance of G1/G0 cells and the absence of neuronal markers such as *elavl3* and *elavl4*. The quiescent-like population clustered separately from the other cells independently of the individual cell cycle phases (as indicated by the transcriptome).

Our analysis not only found several known markers for quiescence in embryonic NPCs, but also shed light on potential novel markers or molecular regulators of embryonic quiescent-like cells. Closer inspection of the genes enriched in the quiescent-like NPC subpopulation indicated consistent enrichment of genes involved in transmembrane transport, cell-cell communication, and mitochondrial metabolism. Differential metabolic states are a key feature distinguishing quiescent stem cell types from their actively proliferating counterparts (Urbán and Cheung, 2021; Velthoven and Rando, 2019). In general, quiescent NSCs depend on glycolysis and fatty acid oxidation (FAO), whereas upon activation, they switch to mitochondrial oxidative phosphorylation and increase lipogenesis (Knobloch and Jessberger, 2017; Llorens-Bobadilla et al., 2015; Urbán and Cheung, 2021). This fits with the higher energetic demands due to increased protein synthesis to support active cell division. Our findings suggest a similar converse metabolic shift upon transition of actively cycling NPCs to slower-cycling quiescent-like embryonic NPCs.

Information regarding the developmental emergence of pre-adult NSCs is sparse in zebrafish. Previous studies of the zebrafish forebrain have demonstrated a significant increase in non-proliferating NPCs between 5 and 7 dpf, which could mean that these NPCs are slow-cycling or pre-quiescent (Alunni et al., 2013; Than-Trong et al., 2018). However, the transcriptional signature accompanying qNSC emergence during zebrafish development was not explored before. Our findings suggest that transcriptionally distinct, quiescent-like embryonic NPCs emerge already at 3 dpf.

In mammalian developing brains, adult NSCs residing in the subventricular zone (V-SVZ NSCs) were shown to stem from a subpopulation of slower-cycling embryonic NPCs (Falk et al., 2017; Fuentealba et al., 2015; Furutachi et al., 2015). This subpopulation emerged at late neurogenic stages and showed transcriptional similarities to adult qNSCs (Yuzwa et al., 2017). Several genes enriched (*mfge8a, cx43, s1pr1, hepacam and aldoc*) in mouse embryonic quiescent-like NPCs were also expressed in the quiescent-like zebrafish NPC subpopulation that we identified. The secreted phagocytosis factor Mfge8a was previously found to be highly enriched in adult qNSC in the mouse hippocampus and is essential for their maintenance (Zhou et al., 2018). The observation that *mfge8a* is, next to adult qNSC in zebrafish and mice (Lange et al., 2020; Yuzwa et al., 2017), also expressed in a subpopulation of embryonic NPCs both in zebrafish and mice suggests that it may play an evolutionary conserved role in the molecular mechanisms that mediate the gradual transition of some embryonic NPCs into the quiescent NSCs of the adult brain.

While our analysis provides evidence of embryonic quiescent-like NPCs, at least in terms of the transcriptome, further evidence regarding their exact identity and biology needs to be obtained. First, it remains to be determined whether these cells are specifically fated to transition into the adult NSCs. To this end, lineage-tracing studies following the full life-cycle of individual NSCs may provide more information. Also, in our analysis, we did not take the regional identity of individual NPCs into account. It is likely that the emergence of pre-quiescent NSCs is not synchronous in the different brain regions. Moreover, local differences in niche architecture and exposure to signaling molecules is likely to contribute to regional differences in qNSC development (Chaker et al., 2016; Dray et al., 2021; Obernier and Alvarez-Buylla, 2019). One intriguing factor here is that adult neurogenesis in zebrafish is much more widespread than in mammals. Therefore, the occurrence of the transition of embryonic into pre-adult NSCs must be widespread in the zebrafish brain. Further exploration of the genes enriched in the quiescent-like population as well as their biology is thus required to obtain more insight into the rules and mechanisms underlying the developmental emergence of pre-adult NSCs.

## Materials and Methods

### Single-cell RNA-seq analysis

#### Computational analysis

The data was analyzed using the Seurat (Satija et al., 2015) v4.3.0 software package for R, v4.2.2 (R Core Team, 2018). Standard read alignment, quality control, normalization, and preliminary rough clustering was adapted from Raj et al. (2020). The resulting gene expression matrices after data preprocessing were loaded into Seurat and analyzed as per Seurat’s proposed clustering workflow. Data was scaled with default parameters when it only included cells in G1/G0. For the merged data containing cells from all cell cycle stages, we removed unwanted variation among cycling cells while maintaining the distinction of cycling and non-cycling cells by regressing out the differences between G2M and S phase as suggested in the Seurat Cell Cycle Scoring and Regression vignette. The top 2000 highly variable genes were used for the further analysis. Dimensionality was first reduced using Principal Component Analysis (PCA), and significant principal components as evaluated by manual inspection of the PC elbow plot were selected for further downstream analysis. We then conducted non-linear dimensionality reduction using Uniform Manifold Approximation Projection (UMAP). Unsupervised clustering was performed using Seurat’s FindNeighbors (with k.param=20) and FindClusters functions. The appropriate resolution used in the FindClusters function for each timepoint was iteratively assessed using the clustree v0.4.4 R package (Zappia and Oshlack, 2018). UMAP visualizations, heatmaps, and dot plots were produced using functions implemented within the Seurat package.

#### Identification and characterization of qNSC-like clusters

Differential gene expression analysis was performed via Seurat’s *FindAllMarkers* function using Wilcoxon rank sum tests. Genes were considered as enriched within a cluster if they were expressed in at least 25% of the cells in a cluster and showed a minimal log2FC of 0.25. To identify which of the clusters could correspond to a population of qNSC-like cells, we tested each cluster’s expression of proposed marker genes for adult qNSCs. We compiled a list of markers used in two studies profiling adult qRGCs in zebrafish (Lange et al. 2020; Than-Trong et al. 2018). We then evaluated which clusters expressed these genes and therefore matched the signature of adult qNSCs. Next, we compared differentially expressed genes in qNSC-like clusters across timepoints to obtain a stable core transcriptional profile of qNSC-like cells over time. Gene ontology analysis was performed using the topGO (Alexa and Rahnenfuhrer, 2017) v2.50.0 R package. GO-terms were considered significantly enriched when reaching an adjusted p-value below 0.05 according to Fisher’s exact test.

### Pseudotime analysis / Pseudotemporal trajectory analysis

Trajectory analysis was performed on the merged dataset including all progenitors using the monocle3 v1.3.1 R package (Trapnell et al., 2014). As the root cluster, we selected the cluster with the most cells from the early stages represented in the dataset. Following trajectory inference, the cells were assigned a pseudotime value along the defined trajectory. Genes with dynamic expression over pseudotime were clustered using k-means clustering based on (Tambalo et al., 2020).

### Workspace specifications

Analysis was performed in RStudio (version 2022.12.0) with R (version 4.2.2). Additional packages used were: clustree_0.4.4, ggraph_2.0.5, KEGGREST_1.34.0, topGO_2.46.0, SparseM_1.8, GO.db_3.14.0, AnnotationDbi_1.56.2, IRanges_2.28.0, S4Vectors_0.32.4, Biobase_2.54.0, graph_1.72.0, BiocGenerics_0.40.0, ggpubr_0.4.0, writexl_1.4.0, xlsx_0.6.5, readxl_1.4.0, viridis_0.6.2, viridisLite_0.4.0, forcats_0.5.1, stringr_1.4.0, dplyr_1.0.9, purrr_0.3.4, readr_2.1.2, tidyr_1.2.0, tibble_3.1.7, tidyverse_1.3.1, ggplot2_3.3.6, sp_1.5-0, SeuratObject_4.1.0, Seurat_4.1.1.

## Code availability

All the code for the data analysis is available on GitHub: github.com/jparidaen.

## Acknowledgements

The authors thank members of the Paridaen lab for the valuable discussions, and in particular Nynke Oosterhof and Glòria Casas Gimeno for advice on single-cell transcriptome analyses.

