## Supplemental information for "Developmental emergence of quiescent-like neural progenitor cells in the zebrafish embryonic brain"

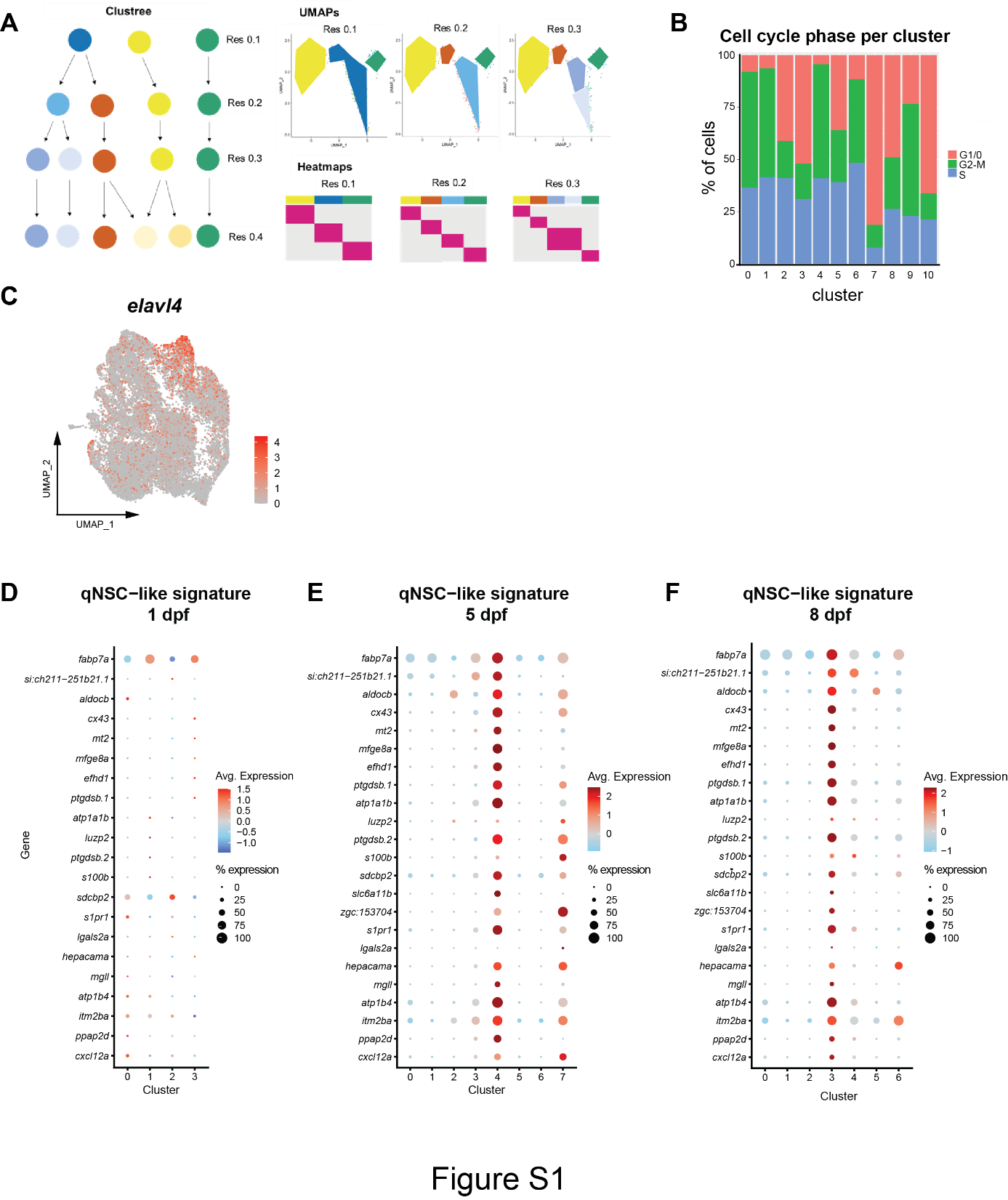

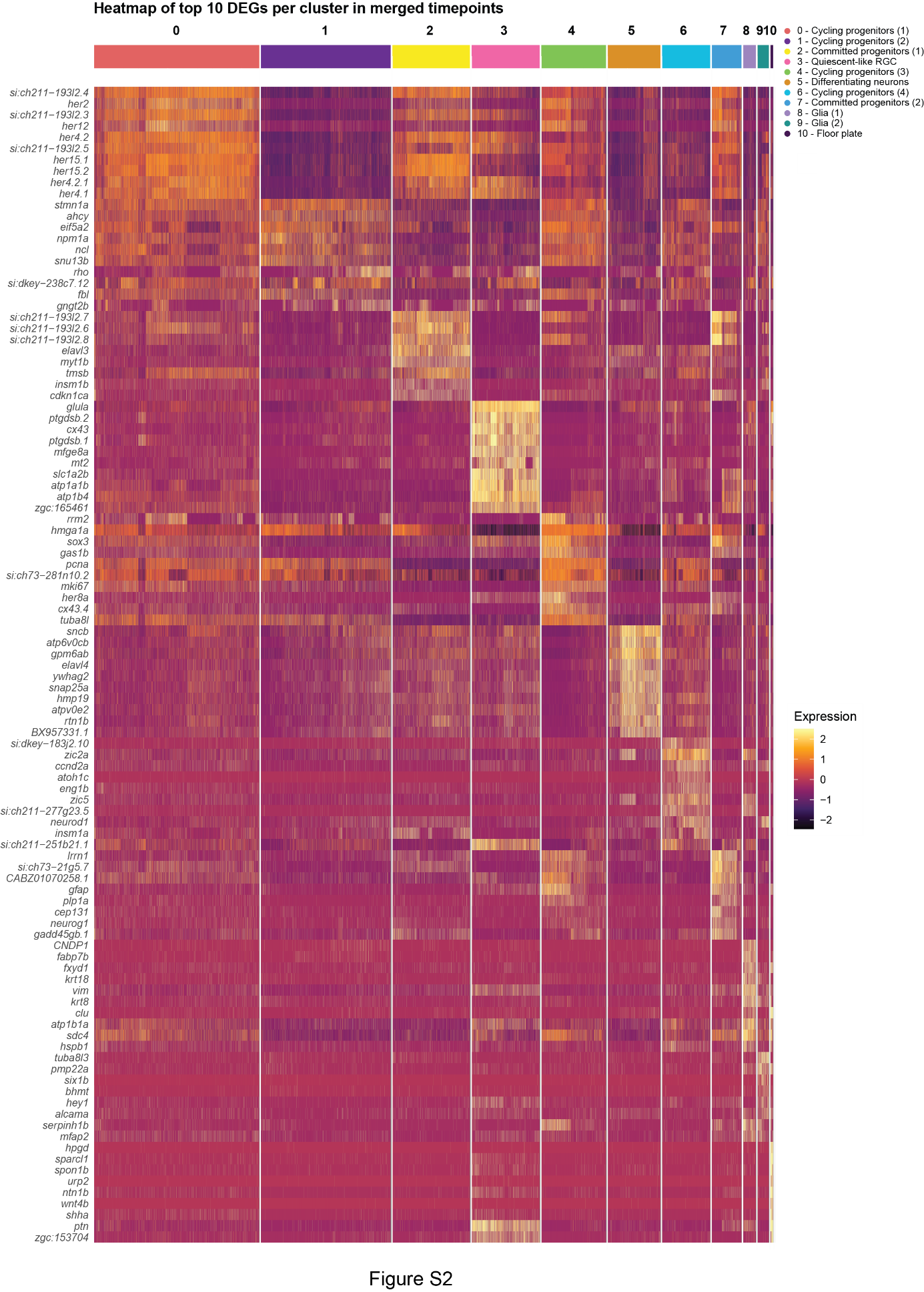

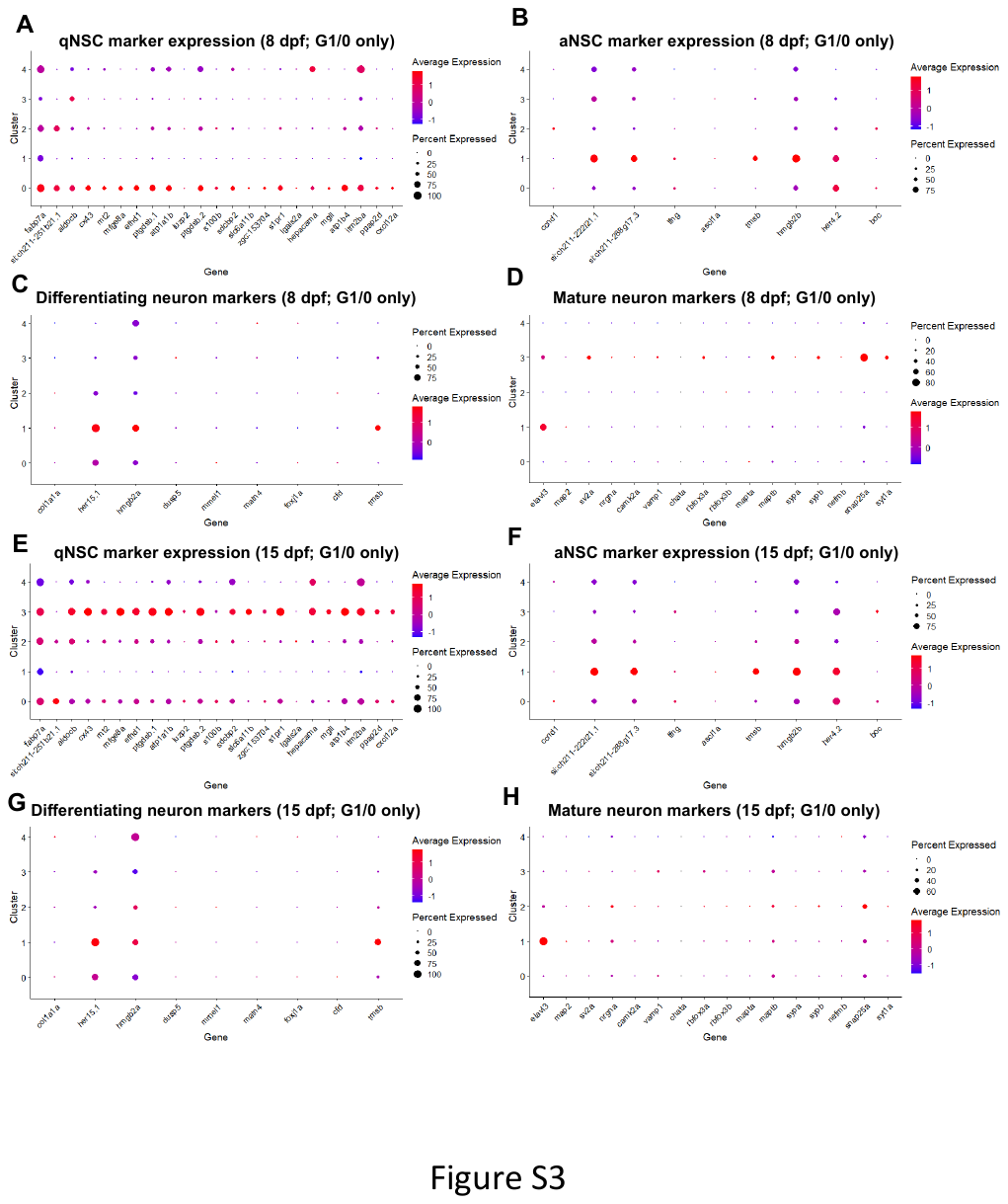

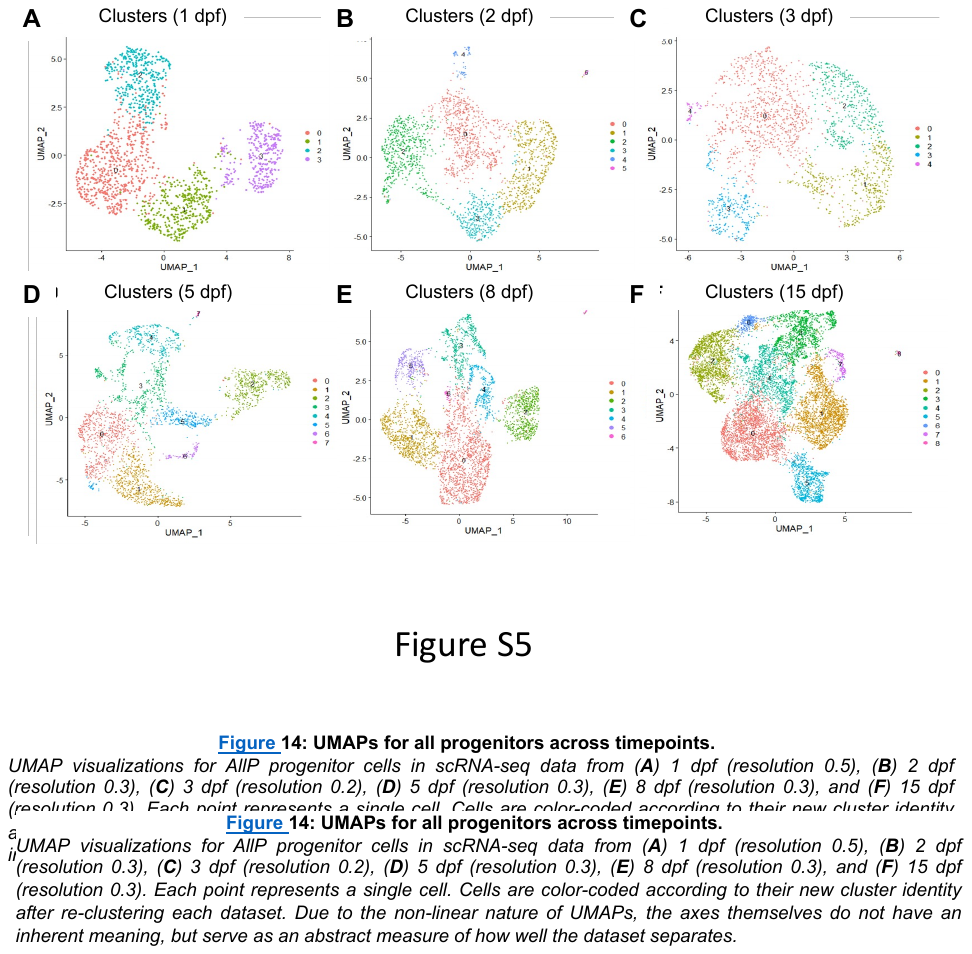

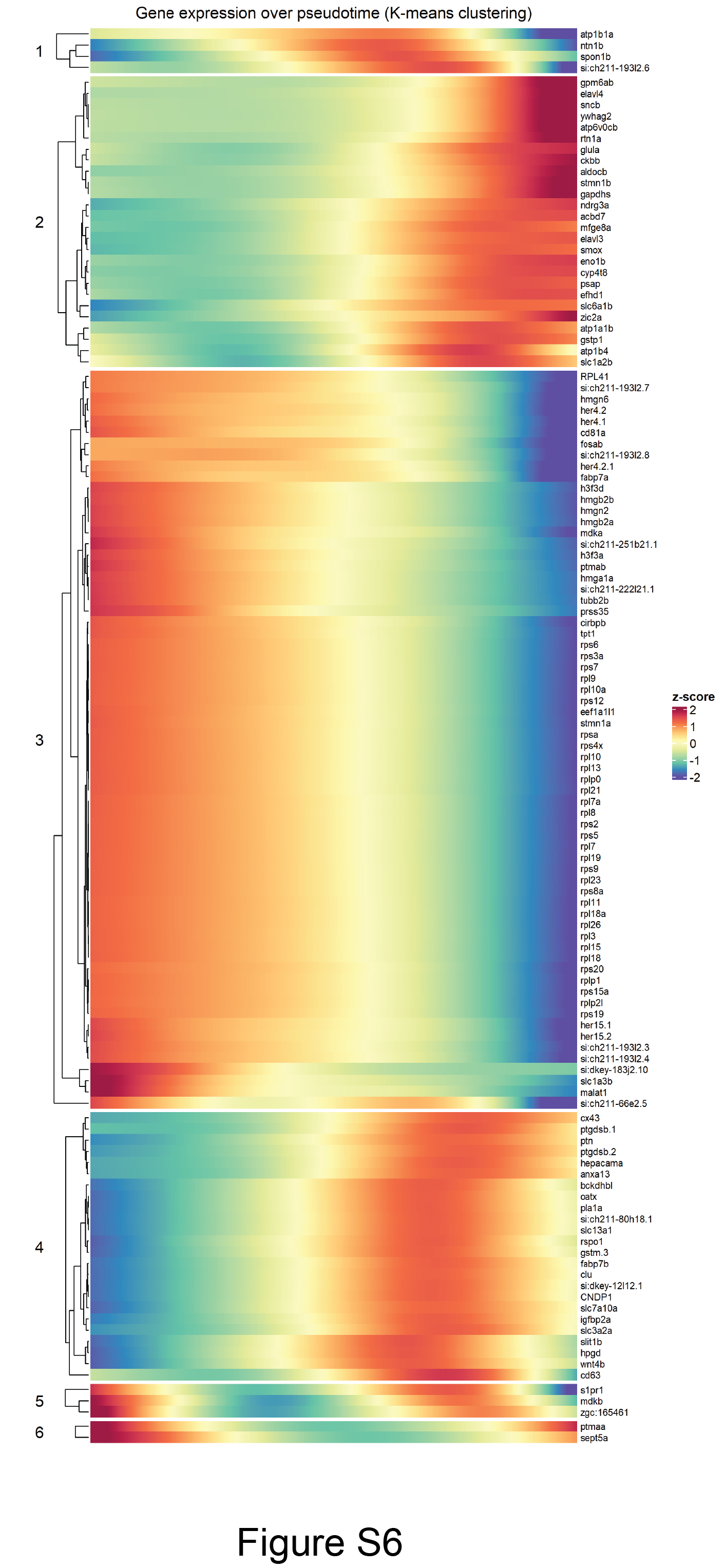

**Supplemental figure legends**

**Figure S1.** **Schematic representation of resolution testing for the *FindClusters* function.**
(A) Clustree plot shows cluster arrangements in multiple resolutions. Resolutions are listed on the right side. Each circle represents one cluster. Arrows represent the origin of cells and to which clusters they are assigned when using a higher resolution. By examining the Clustree for each timepoint, we selected three possible values for the resolution parameter in each dataset. Each value was tested by visualizing the clustering in a UMAP plot (top right) and by generating a heatmap of the top 10 DEGs for each resulting cluster (bottom right). The UMAPs give an overall impression of the rationality of cluster assignments. In this case, the resolution value of 0.2 stands out as the most suitable candidate in this exemplary analysis pipeline. (B) Percentage of individual cells in each cell cycle stage for each subcluster in the all progenitor dataset. (C) UMAP plot (merged dataset) showing the log-normalized expression for the neuronal marker gene *elavl4*. (D-F) Dotplots showing the gene expression of adult qNSC markers at (D) 1 dpf, (E) 5 dpf, and (F) 8 dpf. The y axis shows the individual marker genes and the x axis shows the progenitor clusters at each timepoint. The size of the dots reflects which percentage of cells within a given cluster which express the gene in question while the color of the dots represents the log-normalized expression level with red signifying high expression and blue low expression as shown in the individual keys.

**Figure S2. Gene expression heatmap of the top 10 DEGs in all progenitor clusters within the merged dataset.**
Heatmap for the individual clusters in the merged progenitor datasets. Each column represents one cell arranged according to its assigned cluster as indicated by the colored bars above the figure. Each row corresponds to a gene. Genes are arranged according to the cluster in which they are differentially expressed. The key to the log-normalized expression levels is shown on the right.

**Figure S3. Expression of cell-type specific markers in 8 and 15 dpf G1/0 progenitor clusters.**

Dotplots showing the gene expression of adult qNSC markers at 8 (A) and 15 (E) dpf, adult activated NSC marker genes for 8 (B) and 15 (F) dpf; differentiating neuron marker genes for 8 (C) and 15 (G) dpf, and mature neuron markers for 8 (D) and 15 (H) dpf. The x-axis shows the individual marker genes and the y-axis shows the progenitor clusters at each timepoint. The size of the dots reflects which percentage of cells within a given cluster which express the gene in question while the color of the dots represents the log-normalized expression level with red signifying high expression and blue low expression as shown in the individual keys.

**Figure S4. Clusters and qNSC marker gene expression in the merged all progenitor versus G1/0-only datasets.**

A, C) Unsupervised UMAP plots containing all 22.865 neural progenitors that separate into 11 distinct clusters (A), and 5.218 G1/0-only neural progenitors that separate into 8 distinct clusters (C). (B, D) Projection of the developmental stage of each individual cell onto the UMAP plots for all progenitors (B) and G1/0-only progenitors (D). (E, F) Dotplots showing the gene expression of adult qNSC markers for all neural progenitors (E) and G1/0-only progenitors (F). The x-axis shows the individual marker genes and the y-axis shows the progenitor clusters at each timepoint. The size of the dots reflects which percentage of cells within a given cluster which express the gene in question while the color of the dots represents the log-normalized expression level with red signifying high expression and blue low expression as shown in the individual keys.

**Figure S5. Neural progenitor clusters for each individual developmental timepoint.** Unsupervised UMAP plots containing all neural progenitor cells for each developmental timepoint for 1 dpf (A), 2 dpf (B), 3 dpf (C), 5 dpf (D), 8 dpf (E), and 15 dpf. The distinct cell clusters are indicated with different colors.

**Figure S6. Distinct gene expression dynamics over pseudotime** Heatmap showing all genes with dynamic changes in expression levels over pseudotime. Genes with similar expression dynamics over pseudotime are clustered together using k-means.
